# Complete genome assembly of clinical multidrug resistant *Bacteroides fragilis* isolates enables comprehensive identification of antimicrobial resistance genes and plasmids

**DOI:** 10.1101/633602

**Authors:** Thomas V. Sydenham, Søren Overballe-Petersen, Henrik Hasman, Hannah Wexler, Michael Kemp, Ulrik S. Justesen

**Author notes:** **Corresponding author:** Thomas V. Sydenham.

## Abstract

*Bacteroides fragilis* constitutes a significant part of the normal human gut microbiota and can also act as an opportunistic pathogen. Antimicrobial resistance and the prevalence of antimicrobial resistance genes are increasing, and prediction of antimicrobial susceptibility based on sequence information could support targeted antimicrobial therapy in a clinical setting. Complete identification of insertion sequence (IS) elements carrying promoter sequences upstream of resistance genes is necessary for prediction of antimicrobial resistance. However, *de novo* assemblies from short reads alone are often fractured due to repeat regions and the presence multiple copies of identical IS elements. Identification of plasmids in clinical isolates can aid in the surveillance of the dissemination of antimicrobial resistance and comprehensive sequence databases support microbiome and metagenomic studies. Here we test several short-read, hybrid and long-lead assembly pipelines by assembling the type strain *B. fragilis* CCUG4856T (=ATCC25285=NCTC9343) with Illumina short reads and long reads generated by Oxford Nanopore Technologies (ONT) MinION sequencing. Hybrid assembly with Unicycler, using quality filtered Illumina reads and Filtlong filtered and Canu corrected ONT reads produced the assembly of highest quality. This approach was then applied to six clinical multidrug resistant *B. fragilis* isolates and, with minimal manual finishing of chromosomal assemblies of three isolates, complete, circular assemblies of all isolates were produced. Eleven circular, putative plasmids were identified in the six assemblies of which only three corresponded to a known cultured *Bacteroides* plasmid. Complete IS elements could be identified upstream of antimicrobial resistance genes, however there was not complete correlation between the absence of IS elements and antimicrobial susceptibility. As our knowledge on factors that increase expression of resistance genes in the absence of IS elements is limited, further research is needed prior to implementing antimicrobial resistance prediction for *B. fragilis* from whole genome sequencing.

**REPOSITORIES:** Sequence files (MinION reads de-multiplexed with Deepbinner and basecalled with Albacore in fast5 format and Illumina MiSeq reads in fastq format) and final genome assemblies have been deposited to NCBI/ENA/DDBJ under Bioproject accessions PRJNA525024, PRJNA244942, PRJNA244943, PRJNA244944, PRJNA253771, PRJNA254401, and PRJNA254455

**IMPACT STATEMENT:** Bacterial whole genome sequencing is increasingly used in public health, clinical, and research laboratories for typing, identification of virulence factors, phylogenomics, outbreak investigation and identification of antimicrobial resistance genes. In some settings, diagnostic microbiome amplicon sequencing or metagenomic sequencing directly from clinical samples is already implemented and informs treatment decisions. The prospect of prediction of antimicrobial susceptibility based on resistome identification holds promises for shortening time from sample to report and informing treatment decisions. Databases with comprehensive reference sequences of high quality are a necessity for these purposes. *Bacteroides fragilis* is an important part of the human commensal gut microbiota and is also the most commonly isolated anaerobic bacterium from non-faecal clinical samples but few complete genome assemblies are available through public databases. The fragmented assemblies from short read de novo assembly often negate the identification of insertion sequences upstream of antimicrobial resistance gens, which is necessary for prediction of antimicrobial resistance from whole genome sequencing. Here we test multiple assembly pipelines with short read Illumina data and long read data from Oxford Nanopore Technologies MinION sequencing to select an optimal pipeline for complete genome assembly of *B. fragilis*. However, *B. fragilis* is a highly plastic genome with multiple inversive repeat regions, and complete genome assembly of six clinical multidrug resistant isolates still required minor manual finishing for half the isolates. Complete identification of known insertion sequences and resistance genes was possible from the complete genome. In addition, the current catalogue of *Bacteroides* plasmid sequences is augmented by eight new plasmid sequences that do not have corresponding, complete entries in the NCBI database. This work almost doubles the number of publicly available complete, finished chromosomal and plasmid *B. fragilis* sequences paving the way for further studies on antimicrobial resistance prediction and increased quality of microbiome and metagenomic studies.

**DATA SUMMARY:**

1. Sequence read files (Oxford Nanopore (ONT) fast5 files and Illumina fastq files) as well as the final genome assemblies have been deposited to NCBI/ENA/DDBJ under Bioproject accessions PRJNA525024, PRJNA244942, PRJNA244943, PRJNA244944, PRJNA253771, PRJNA254401, and PRJNA254455.
2. Fastq format of demultiplexed ONT reads trimmed of adapters and barcode sequences are available at doi.org/10.5281/zenodo.2677927
3. Genome assemblies from the assembly pipeline validation are available at doi: doi.org/10.5281/zenodo.2648546.
4. Genome assemblies corresponding to each stage of the process of the assembly are available at doi.org/10.5281/zenodo.2661704.
5. Full commands and scripts used are available from GitHub: https://github.com/thsyd/bfassembly as well as a static version at doi.org/10.5281/zenodo.2683511

## INTRODUCTION

*Bacteroides fragilis* is a Gram-negative anaerobic bacterium that is commensal to the human gut but can act as an opportunistic pathogen; it is the most commonly isolated anaerobic bacteria from non-faecal clinical samples (1). Antimicrobial resistance rates are increasing for *B. fragilis*, especially for carbapenems and metronidazole, two widely used antimicrobials for treatment of severe infections and anaerobe bacteria (2,3). Antimicrobial susceptibility testing of anaerobes using agar dilution or gradient strip methods can be costly and labour intensive and despite efforts to validate disk diffusion as a less expensive option, turn-around time will still be least 18 hours and validation for individual species will be required (4).

Antimicrobial resistance prediction from bacterial whole genome sequences, from cultured isolates as well as metagenomes, could be implemented in clinical microbiology in the near future, with the potential for improved sample-to-report turnover time and possibly eliminating the need for phenotypical testing for individual species (5–8). For a few species, prediction of antimicrobial resistance from WGS has been validated, but for the majority of clinical relevant species challenges still remain (6,9,10).

Based on DNA-DNA hybridisation studies, *B. fragilis* can be divided into two DNA homology groups (division I and II), whose ribosomal contents are so different that the two divisions can be distinguished by mass spectrometry routinely used to identify isolates in clinical laboratories (11). *B. fragilis* division I carry the chromosomal cephalosporinase gene *cepA* whilst *B. fragilis* division II harbour the chromosomal metallo-β-lactamase gene *cfiA* (also known as *ccrA*) (12,13). The *cfiA* gene can confer resistance to carbapenems, a class of antimicrobials usually reserved for patients with severe sepsis or infections with multidrug-resistant bacteria. But expression levels are partly controlled by insertion sequence (IS) elements carrying promotor sequences inserted upstream of the gene and only 30-50% of clinical isolates that harbour *cfiA* display phenotypically reduced susceptibility to carbapenems (3). The same pattern of expression control can be observed for genes associated with resistance to metronidazole (*nim* genes) and clindamycin (*erm* genes) (1).

In 2014 we observed that identification of IS elements upstream of known antimicrobial resistance genes in *B. fragilis* was hampered in short read *de novo* assemblies even though the genes could be identified (14). This occurred because contigs were often terminated close to the start of the resistance genes, presumably due to the proliferation of multiple copies of the same IS elements throughout the *B. fragilis* genomes. Genome assemblies from short read sequencing technologies alone most often result in fragmented assemblies because of repetitive regions and genome elements with multiple occurrences in the chromosomes and plasmids (15,16). Therefore, we could not predict antimicrobial resistance (AMR) phenotypes in *B. fragilis* using only short reads for WGS since IS element identification is a prerequisite for correct genotype-phenotype associations. Long read sequencing technologies are increasingly being utilised to increase the contiguity of bacterial genome assemblies and often result in complete, closed chromosomes and plasmids (17–20). This provides possibilities for comprehensive identification of IS elements, insights into genome structures and characterisation of other mobilisable elements and associated genes. Complete identification and characterisation of plasmids in sequenced isolates would allow for improved analysis of the plasmid-mediated spread of antimicrobial resistance.

Bioinformatic analysis of WGS data depends heavily on high-quality reference databases. Anaerobes make up most of the bacterial human commensal microbiota but are most likely underrepresented in public databases of whole genomes from cultured isolates. The NCBI Genome database (accessed 31-03-2019) contains genome sequences of 191,411 bacteria of which 13,483 are marked as complete assemblies. Only seven of these are *Bacteroides fragilis* (21–27). In comparison there are 776 assemblies of *E. coli* marked as complete and 398 of *S. aureus*. Improving the representation of complete assemblies of *B. fragilis* in the public genome databases will support the development of antimicrobial resistance prediction from WGS as well as microbiome and metagenomic analysis projects.

The aims of this study were to select an optimal assembly software pipeline for complete, circular assembly of *Bacteroides fragilis* and demonstrate the utility of complete assembly for both plasmid identification and comprehensive detection of genes and IS elements associated with antimicrobial resistance. We assembled the *B. fragilis* CCUG4856T (= ATCC25285 = NCTC9343) reference strain utilising long reads generated with the MinION sequencer from Oxford Nanopore Technology (ONT) and high-quality Illumina short reads and selected the best assembly pipeline by comparing assemblies to the Sanger sequenced reference NCTC9343 (RefSeq accesion GCF_000025985.1). The best assembly pipeline was then applied to six clinical multi-drug resistant *B. fragilis* isolates from our 2014 study (14).

## METHODS

### Culture conditions and DNA extraction

*Bacteroides fragilis* CCUG4856T and the six strains described in our previous study were included (14,21). Strains were stored at −80° in beef extract broth with 10% glycerol (SSI Diagnostica) and cultured on solid chocolate agar with added vitamin K and cysteine (SSI Diagnostica) for 48 hrs in an anaerobic atmosphere at 35 °C. Ten µl of culture was transferred to 14 ml saccharose serum broth (SSI Diagnostica) and incubated for 18 hrs under the same conditions. DNA was then extracted using the Genomic-Tip G/500 kit (Qiagen) following the manufacturers protocol for Gram negative bacteria and eluted into 5 mM Tris pH 7.5 0.5 mM EDTA buffer. Quality control was performed by measuring fragment length on a TapeStation 2500 (Genomic DNA ScreenTape, Agilent), purity on the NanoDrop (ThermoFisher Scientific) and concentration on the Qubit (dsDNA BR kit; Invitrogen). The eluted DNA was then stored at −20 °C.

### Illumina library preparation, sequencing and quality control

The strains had previously been sequenced and assembled using Illumina short reads for our previous study (14), but to minimise biological disparities we opted to re-sequence with Illumina using the same DNA extraction prepared for long read sequencing. Paired-end libraries were generated using the Nextera XT DNA sample preparation kit (Illumina) according to the manufacturer’s protocol. DNA was sequenced on a MiSeq sequencer (Illumina) with 150 bp reads for a theoretical read depth of 100x. Read quality metrics were evaluated using FastQC (https://www.bioinformatics.babraham.ac.uk/projects/fastqc/) and fastp v0.19.6 (28). Filterbytile from the BBmap package (http://sourceforge.net/projects/bbmap/) was used for removing low-quality reads based on positional information on the sequencing flowcell and TrimGalore (http://www.bioinformatics.babraham.ac.uk/projects/trim_galore/), with settings --qual 20 and --length 126, provided additional adapter and quality trimming. FastQ files were then randomly down-sampled to < 100x crude read depth using an estimated genome size of 5.3 Mb, as higher read depths tend to reduce assembly quality (29).

### Nanopore library preparation and MinION sequencing

Sequencing libraries were prepared using the Rapid Barcoding kit (SQK-RPB004; Oxford Nanopore Technologies) following the manufacturers protocol (version RPB_9059_v1_revC_08Mar2018) with SPRI bead clean up (AMPure XT beads; Beckman Coulter) as described. Sequencing was performed as multiplex runs on a MinION connected to a Windows PC with MinKnow v1.15.1 using FLO-MIN106 R9.4 flowcells. Raw fast5 files were transferred to the Computerome high performance cluster (https://www.computerome.dk/) for analysis. Four sequencing runs were performed, as the first two runs did not provide enough data for complete assembly of all isolates (see results section).

### Fast5 demultiplexing, base-calling, quality control and filtering

The raw fast5 files were demultiplexed with Deepbinner v0.2.0 and base-called using Albacore v2.3.3, retaining only those barcodes Deepbinner and Albacore agreed upon for minimal barcode misclassification (30). Porechop v0.2.4 (https://github.com/rrwick/Porechop) with the *--discard_middle* option was used for adapter and barcode trimming and read statistics were collected using NanoPlot (31). Filtlong v0.2.0 (https://github.com/rrwick/Filtlong) was used to filter the long reads by either removing the worst 10% or by retaining 500Mbs in total, which ever option resulted in fewer reads.

### Assembly validation

To select and validate the optimum assembly pipeline *Bacteroides fragilis* CCUG4856T was assembled using a variety of well-known assemblers and polishing tools (Table 1). Each assembler was run with the Filtlong filtered reads as input or the filtered reads corrected with Canu 1.8 (with standard settings, corMinCoverage=0, or coroutCoverage=999). Canu was also tested with the unfiltered reads as input. Hybrid assemblers used the filtered long reads and the filtered, trimmed and down-sampled Illumina reads. Unicycler includes polishing with Racon and Pilon. For assemblers other than Unicycler, Racon polishing with ONT reads was run for one or two rounds and Pilon was run until no changes were made or for a maximum of six rounds. Racon polishing with Illumina reads was run for one round.

**Table 1.** Genome assemblers and polishing tools tested.

| Genome assembler and version | Link | Reference |
| --- | --- | --- |
| Wtdbg2 v2.3 | https://github.com/ruanjue/wtdbg2 | (70) |
| Miniasm v0.3r179 | https://github.com/lh3/miniasm | (39,71) |
| Flye v2.3.7 | https://github.com/fenderglass/Flye | (59,72) |
| Canu v1.8 | https://github.com/marbl/canu | (73) |
| Spades (including Hybridspades) v3.13.0 | https://github.com/ablab/spades | (74,75) |
| Skesa v2.3.0 | https://github.com/ncbi/SKESA | (76) |
| Unicycler v0.4.7 | https://github.com/rrwick/Unicycler | (77) |
| **Assembly polishing tools** |  |  |
| Nanopolish v0.10.2 | https://github.com/jts/nanopolish | (78) |
| Racon v1.3.1 | https://github.com/isovic/racon | (79) |
| Pilon v1.22 | https://github.com/broadinstitute/pilon | (80) |

The original Sanger sequenced *Bacteroides fragilis* NCTC9343 (=CCUG4856T) (21) downloaded from NCBI RefSeq (accession GCF_000025985.1) was used as reference sequence for the assembly comparisons and Quast v5.0.2 was used for assembly summary statistics, indel count, and K-mer-based completion (32). BUSCO v3.0.2b with the bacteroidetes_odb9 dataset, CheckM v1.0.12, and Prokka v1.13.3 were used to assess gene content (33–35). Average nucleotide identity was calculated using https://github.com/chjp/ANI/blob/master/ANI.pl and ALE v0.9, which uses a likelihood based approach to assess the quality of different assemblies, was also used to score the assemblies (36,37). Ranking of assemblies was based on number of contigs, number of circular contigs, closeness to total length compared to the reference genome, number of local misassembles, number of mismatches per 100 kb, number of indels per 100kb, average nucleotide identity (ANI), CheckM and BUSCO scores, and the total ALE score (a higher score is better). Please see https://github.com/thsyd/bfassembly for full bioinformatics methods.

### Genome assembly of MDR *B. fragilis* isolates

The assembly strategy deemed to produce the highest quality genome for CCUG4856T was chosen for initial assembly of the six MDR *B. fragilis* isolates. Manual finishing of incomplete assemblies was performed using Bandage for visualisation of assembly graphs and BLASTn searches (38). Minimap2 and BWA MEM were used to map reads to the assemblies for coverage graphs (39,40). Long read assembly with Flye was compared to the Unicycler assembly and used to guide and validate the manual finishing results. Circlator’s *fixstart* task was used to fix the start position of the manually finished genomes to be at the *dnaA* gene (41).

The assembled genomes were submitted to NCBI GenBank and annotated with PGAP (42). ABRicate v0.8.10 (https://github.com/tseemann/ABRicate) (with options --minid 40 --mincov 25) was used to screen for antimicrobial resistance genes with the ResFinder (database date 19-08-2018), NCBI Bacterial Antimicrobial Resistance Reference Gene Database (database date 19-09-2018), and CARD (v2.0.3) databases, supplemented with nucleotide sequences for the multidrug efflux-pump genes *bexA* (GenBank: AB067769.1:3564..4895) and *bexB* (GenBank: AY375536.1:4599..5963) (43,44). IS elements were identified using ABRicate with data from the IS-finder database (http://www-is.biotoul.fr/, Update: 2018-07-25) (45).

### Identification of plasmids and mobile genetic elements

The PLSDB web server (https://ccb-microbe.cs.uni-saarland.de/plsdb/) (data v. 2019_03_05) contains bacterial plasmid sequences retrieved from the NCBI and was used for screening and identifying putative plasmids sequences (46). Only hits to accessions from cultured organisms were included. Putative plasmids not identified using PLSDB, were evaluated by the read depth relative to the chromosome (higher relative read depth indicates plasmid sequence) and Pfam families covering known plasmid replication domains from Table 1 in reference (47) were downloaded from the Pfam database (Pfam 32.0, https://pfam.xfam.org/) and used for screening putative plasmids with ABRicate.

## RESULTS

### Sequencing data quality

For Illumina data, a median of 3,465,082 reads (interquartile range [IQR]: 3,177,493-5,001,077) were generated for each isolate (Supplementary Table S1)). After filtering, adapter-removal and down sampling a median of 449,022,741 bases (IQR: 433,517,549-530,257,210) were available per isolate with 87-96% Q30 bases corresponding to calculated read depths of 75-103%. The %GC content of the reads for each isolate (median 42.9%, range: 42.6-43.3%) were very consistent and within the expected range for the *Bacteroides* genus (40-48%) (48).

Isolates were sequenced in runs multiplexed with other isolates not included in this study. Based on initial test assemblies using Unicycler without filtering or Canu correction (not shown) it was concluded that data from the first ONT sequencing runs were to be supplemented by additional runs to increase the chance of complete assembly of all isolates. Concatenating reads from runs, a median of 75,598 reads [IQR: 50,210-112,065] with a median length of 2,938-4,393 bases were generated for each isolate (Supplementary Table S1). Filtering with Filtlong and correction with Canu resulted in a median of 8,515 reads (IQR: 6,226-10,370) with median lengths of 6,181-38,588 for each isolate as input for the assemblies.

### Selecting the optimal assembly pipeline

141 assemblies of *B. fragilis* CCUG4856T were generated using the various assemblers and polishing steps (Supplementary Table S2). Compared to the reference genome, Unicycler assemblies were of the highest quality (**Table 2**). Unicycler, with any of the read input options, produced two circular contigs of the expected lengths, and the differences between the various Unicycler assemblies were minimal (**Table 3**). Assemblies with Canu corrected reads showed slightly higher genome fractions and average nucleotide identities to the reference and fewer mismatches and indels, when compared to Unicycler alone. Unicycler assemblies corrected with Racon using Illumina reads worsened slightly overall with 0.04-0.19 more indels and 0.14-0.25 more mismatches per 100 kbp. Based on this initial evaluation, the assembly pipeline using Canu corrected reads with default options was chosen (Assembly “OF.CS” in **Table 3**). This would reduce the number of long reads, compared to Canu correction with corMinCoverage=0, or coroutCoverage=999, and thereby lead to a faster run-time for Unicycler.

**Table 2.** Selected quality indicators for the best genome assembly of *B. fragilis* CCUG4856T per assembly pipeline. RefSeq accession GCF_000025985.1 was used as reference. OF: ONT reads filtered with Filtlong, CS: Canu corrected standard settings, CM: Canu corrected with option corMinCoverage=0, CO: Canu corrected with option coroutCoverage=999, RO2: Two rounds of Racon polishing with ONT reads, RI: Racon polishing with Illumina reads, PI[n]: Pilon polishing with Illumina reads, [n] rounds. Full results are available in Supplementary Table S2.

| Assembly | Contigs | Largest contig | Total length | Mis-assemblies | Genome fraction (%) | Mismatches per 100 kbp | Indels per 100 kbp | Average nucleotide identity | CheckM Completeness | BUSCOs: complete and single-copy/complete and duplicate/fragment (of 443) | Prokka genes | Prokka rRNA | Prokka tRNA | Total ALE score |
| --- | --- | --- | --- | --- | --- | --- | --- | --- | --- | --- | --- | --- | --- | --- |
| GCF_000025985.1 | 2 | 5,205,140 | 5,241,700 | 0 | 100.000 | 0 | 0 | 100.0000 | 99.26 | 442/0/1 | 4439 | 19 | 73 | -17071758.95 |
| Skesa | 46 | 553,341 | 5,201,945 | 3 | 99.237 | 0.23 | 0.15 | 99.9980 | 99.26 | 440/2/1 | 4391 | 2 | 62 | -20926329.69 |
| Spades | 23 | 1,779,941 | 5,212,217 | 4 | 99.396 | 0.44 | 0.17 | 99.9866 | 99.26 | 440/2/1 | 4407 | 3 | 56 | -19676529.39 |
| Canu.OF.CO.RO2.RI.PI3 | 2 | 5,247,938 | 5,350,432 | 8 | 99.972 | 4.94 | 15.9 | 99.9752 | 99.26 | 442/0/1 | 4634 | 19 | 73 | -19283611.73 |
| Flye.OF.CS.PI5.RI | 5 | 2,282,650 | 5,269,269 | 4 | 99.917 | 1.07 | 6.24 | 99.9781 | 99.26 | 441/1/1 | 4476 | 19 | 73 | -18222322.23 |
| Miniasm.OF.CM.RO2.PI5 | 3 | 5,204,445 | 5,277,434 | 2 | 99.972 | 5.21 | 17.75 | 99.9691 | 98.88 | 442/0/1 | 4607 | 19 | 73 | -17789234.97 |
| Wtdbg2.OF.CO.RO2.PI6.RI | 3 | 5,192,352 | 5,234,448 | 7 | 99.723 | 3.23 | 3.04 | 99.9807 | 99.26 | 442/0/1 | 4437 | 19 | 73 | -18750266.21 |
| SpadesHybrid.CS | 5 | 3,093,122 | 5,242,724 | 7 | 99.987 | 1.89 | 0.53 | 99.9856 | 99.26 | 440/2/1 | 4441 | 19 | 73 | -18535980.68 |
| Unicycler.OF.CS | 2 | 5,205,133 | 5,241,693 | 2 | 99.972 | 0.84 | 0.48 | 99.9997 | 99.26 | 442/0/1 | 4435 | 19 | 73 | -17200232.52 |

**Table 3.**
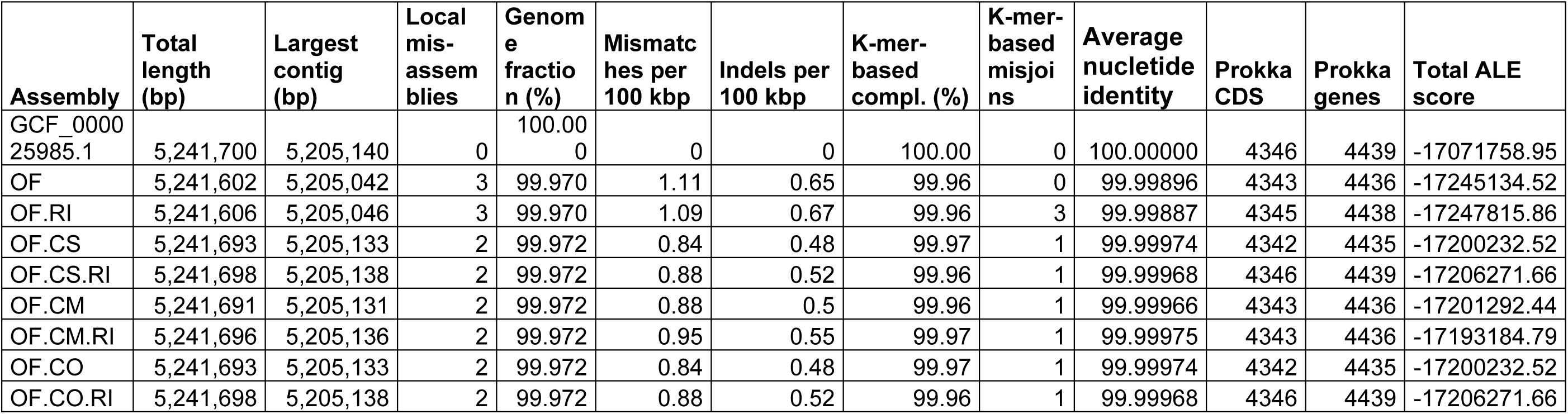
Hybrid Unicycler assemblies of *B. fragilis* CCUG4856T. RefSeq accession GCF_000025985.1 was used as reference. OF: ONT reads filtered with Filtlong; CS: Canu corrected standard settings; CM: Canu corrected with option corMinCoverage=0; CO: Canu corrected with option coroutCoverage=999; RI: Racon polishing with Illumina reads. Unicycler performs assembly polishing with Racon (ONT reads) and Pilon. Full results are available in Supplementary Table S2.

| Assembly | Total length (bp) | Largest contig (bp) | Local mis-assemblies | Genome fraction (%) | Mismatches per 100 kbp | Indels per 100 kbp | K-mer-based compl. (%) | K-mer-based misjoins | Average nucleotide identity | Prokka CDS | Prokka genes | Total ALE score |
| --- | --- | --- | --- | --- | --- | --- | --- | --- | --- | --- | --- | --- |
| GCF_000025985.1 | 5,241,700 | 5,205,140 | 0 | 100.000 | 0 | 0 | 100.00 | 0 | 100.00000 | 4346 | 4439 | -17071758.95 |
| OF | 5,241,602 | 5,205,042 | 3 | 99.970 | 1.11 | 0.65 | 99.96 | 0 | 99.99896 | 4343 | 4436 | -17245134.52 |
| OF.RI | 5,241,606 | 5,205,046 | 3 | 99.970 | 1.09 | 0.67 | 99.96 | 3 | 99.99887 | 4345 | 4438 | -17247815.86 |
| OF.CS | 5,241,693 | 5,205,133 | 2 | 99.972 | 0.84 | 0.48 | 99.97 | 1 | 99.99974 | 4342 | 4435 | -17200232.52 |
| OF.CS.RI | 5,241,698 | 5,205,138 | 2 | 99.972 | 0.88 | 0.52 | 99.96 | 1 | 99.99968 | 4346 | 4439 | -17206271.66 |
| OF.CM | 5,241,691 | 5,205,131 | 2 | 99.972 | 0.88 | 0.5 | 99.96 | 1 | 99.99966 | 4343 | 4436 | -17201292.44 |
| OF.CM.RI | 5,241,696 | 5,205,136 | 2 | 99.972 | 0.95 | 0.55 | 99.97 | 1 | 99.99975 | 4343 | 4436 | -17193184.79 |
| OF.CO | 5,241,693 | 5,205,133 | 2 | 99.972 | 0.84 | 0.48 | 99.97 | 1 | 99.99974 | 4342 | 4435 | -17200232.52 |
| OF.CO.RI | 5,241,698 | 5,205,138 | 2 | 99.972 | 0.88 | 0.52 | 99.96 | 1 | 99.99968 | 4346 | 4439 | -17206271.66 |

The hybrid Unicycler assembly of CCUG4856T with standard Canu corrected ONT reads consists of two circular contigs of 5,205,133 and 36,560 bp in length. The plasmid is the same length as plasmid pBF9343 from the reference assembly GCF_000025985.1 and the chromosome is seven bases shorter. Alignments of the Sanger sequenced assembly GCF_000025985.1 with the hybrid Unicycler assembly show an 88,045 bp inversion in the hybrid assembly compared to the Sanger assembly (Figure 1). This inversion is present in all the best assemblies, including assemblies derived from solely ONT sequences or Illumina sequences (Supplementary Figure S1) as well as two additional assemblies of NCTC9343/ATCC25285 from PacBio and Illumina sequences downloaded from NCBI RefSeq (Supplementary Figure S2).

**Figure 1.**
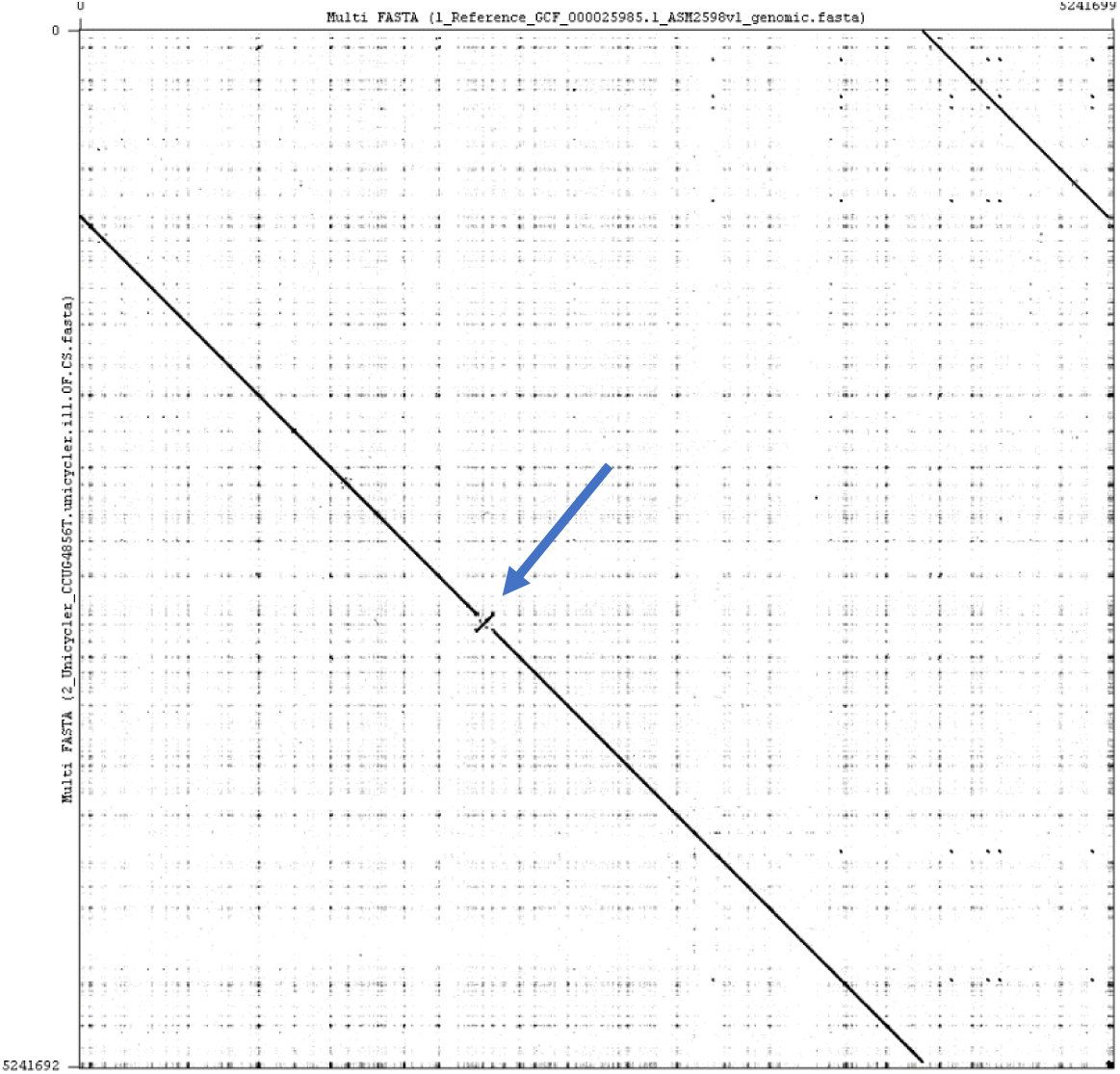
Dot plot matrix of the alignment of the reference assembly and the hybrid Unicycler assembly using Gepard v1.40 (81). The *B. fragilis* NCTC9343 (RefSeq GCF_000025985.1) reference assembly derived from Sanger sequencing is on the x-axis and the hybrid Unicycler assembly on the y-axis. On this otherwise near perfect alignment with high similarity, an 88,045 bp inversion with 100% ID is observed at nucleotide positions 2,941,962..3,030,006 on the Unicycler assembly (2,005,742..2,093,786 on the reference sequence) (indicated by the blue arrow).

### Complete assembly of six multidrug resistant isolates

Unicycler, using filtered and trimmed Illumina reads and the Filtlong filtered and Canu corrected ONT reads from the first sequencing runs, generated complete, continuous, circular assemblies for two of the six isolates (BFO18 and BFO67) (Figure 2). For the assemblies that were not complete with sequencing data from the first MinION runs, increasing the amount of ONT data resulted in fewer contigs overall, except for BF067, where the additional data from the second sequencing run led to a fragmented assembly and manual finishing was necessary. Performing assembly of isolate S01 without Canu correction of the ONT reads from the first sequencing resulted in a closed chromosome and performing Canu correction of reads resulted in a fragmentation of the chromosome. This was ameliorated by including more ONT data. By manual finishing using read mapping and additional assembly with Flye, the remaining three assemblies were circularised. Chromosomes varied in length from 5,141,257 – 5,504,076 bp. Alignment of ONT and Illumina reads to the chromosome assemblies showed even coverage for both sequencing technologies (Supplementary Figure S3). For BFO85 a >100% relative read depth increase was observed at approximately 25kb-38kb. This could represent a 12 kb repeat region that was not resolved in the assembly. Seven (47%) of the 15 PGAP annotated CDS’ in the 13kb region were annotated as hypothetical proteins. None of the annotated CDS’ represented mobilisable proteins.

**Figure 2.**
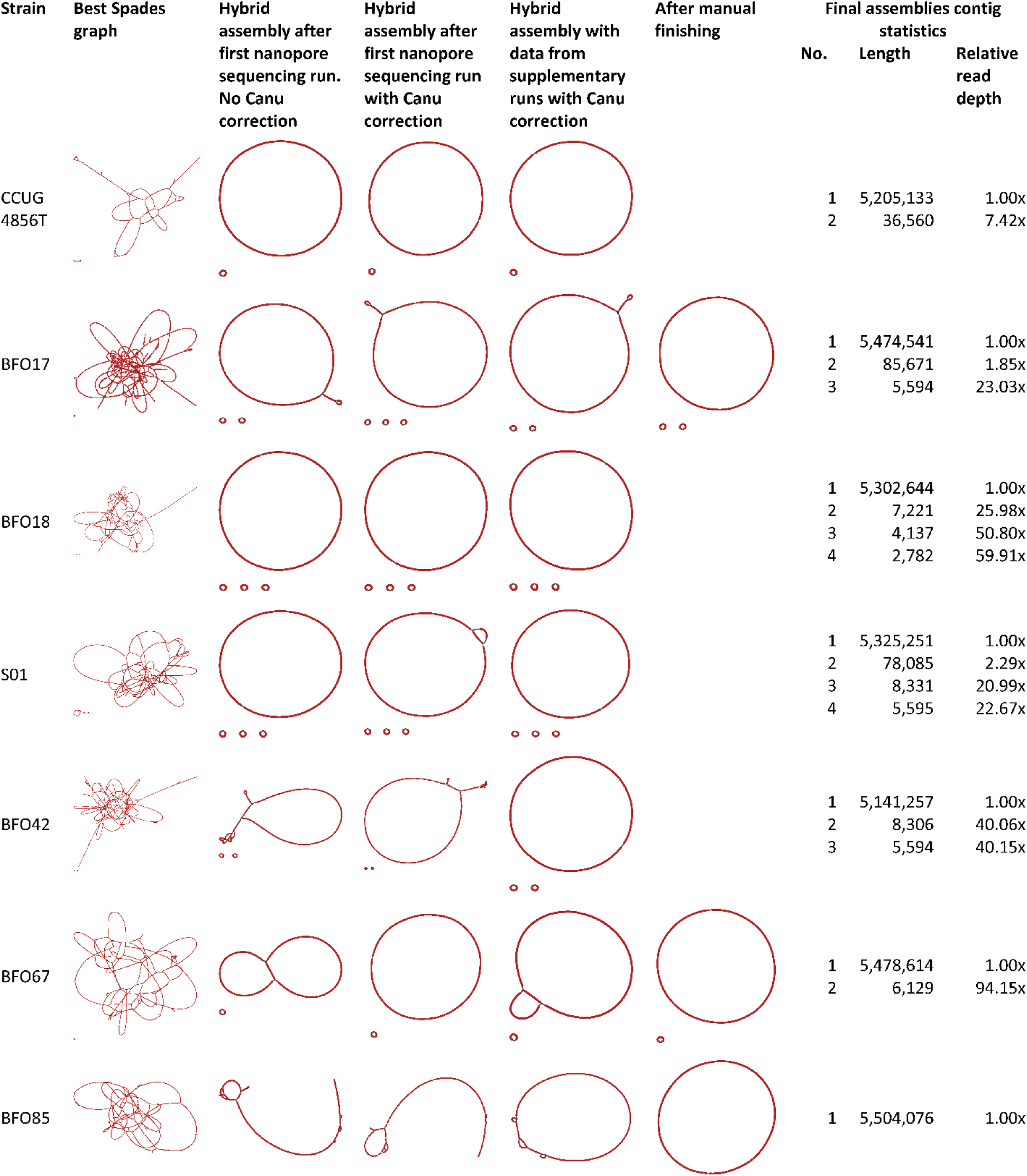
Evolution of genome assemblies with added data and manual finishing. The best SPAdes assembly graphs by Unicycler with short reads only are shown on the far left. Supplying ONT reads improved the assemblies overall, but only three were circularised with singular chromosome contigs with data from the initial MinION sequencing runs. Adding additional ONT data and correcting reads with Canu did not improve assemblies for all isolates. Manual finishing was necessary to finish assemblies for three isolates. Assembly graph images generated with Bandage. Read information can be found in Supplementary Table S1.

### Eleven putative plasmid sequences were identified

A total of 11 putative circular plasmids were identified in the six *B. fragilis* isolates (**Table 4**). Zero to three putative plasmids were identified per isolate with lengths varying from 2,782 to 85,671bp.

**Table 4.**
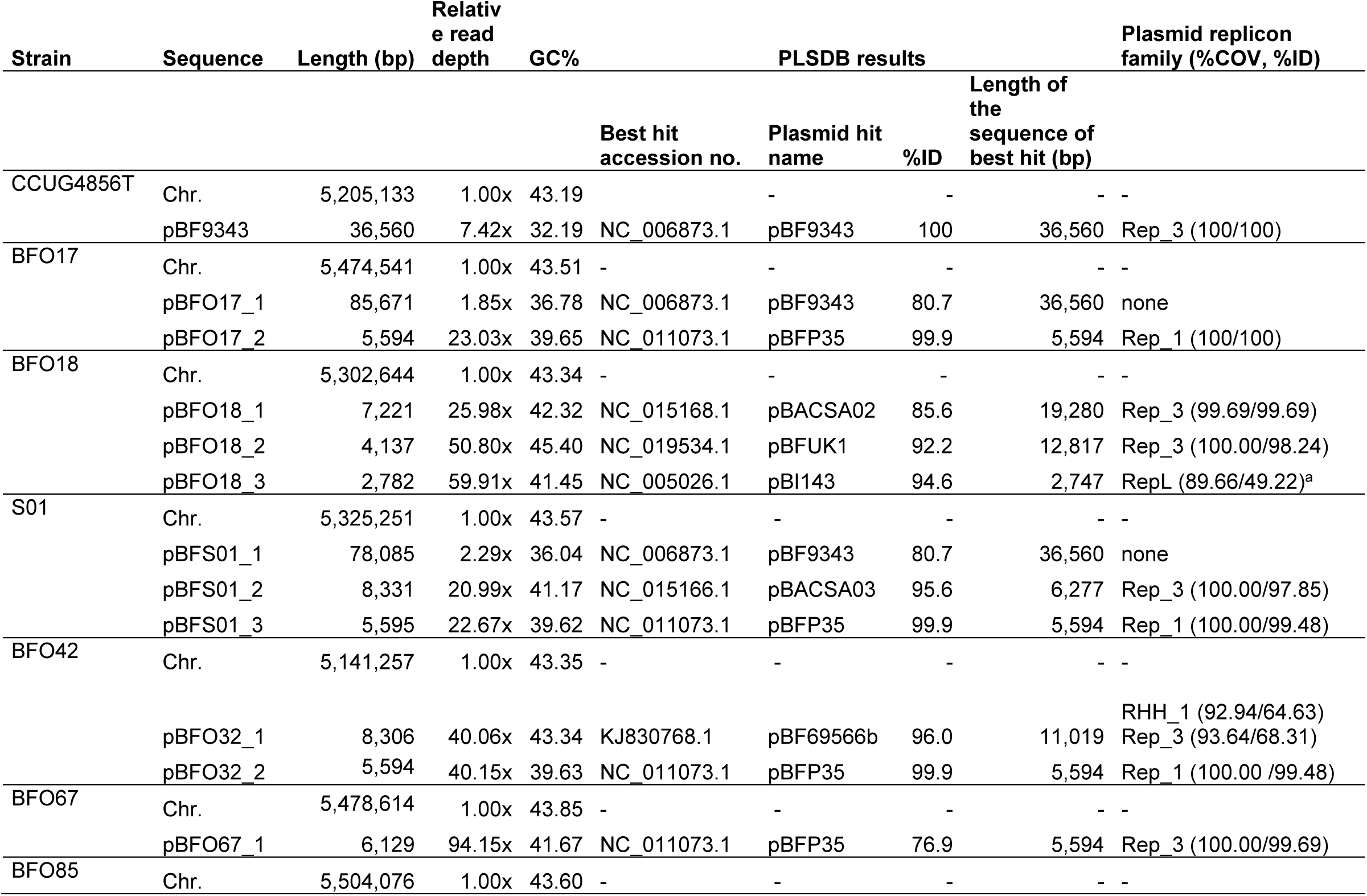
Putative plasmid sequences of the complete *B. fragilis* assemblies. Putative plasmid sequences from the hybrid assemblies of *B. fragilis* CCUG4856T and the six MDR *B. fragilis* isolates were screened using the PLSDB. The best hit to plasmids from cultured isolates is shown. Only three putative plasmids from the MDR *B. fragilis* isolate assemblies could be identified with confident %ID. For most sequences, plasmid replication family proteins were identified in the putative plasmids using ABRicate with a database of sequences downloaded from the Pfam database, strengthening the interpretation that the circularised putative plasmid sequences do in fact represent plasmids harboured by the isolates. Notes: ^a^Annotated as RepA protein in the PGAP annotation. Abbreviations: %ID, %COV, no.; number, Chr.; chromosome.

The PLSDB database contains NCBI RefSeq plasmid sequences marked as complete. Three of the 11 putative plasmid sequences were found to match (ID > 98%) a sequence in PLSDB (Table 4). These three all matched the cryptic plasmid pBFP35 (49). The NCBI Nucleotide database was queried using BLASTn with the remaining unidentified putative plasmid sequences (50). BFO18 putative plasmid sequence pBFO18_1 (7,221 bp) resembles plasmid pIP421, a 7.2kb plasmid with metronidazole resistance gene *nimD* and IS*1169*. Partial sequences in NCBI GenBank spanning the *nimD* gene, IS element and *RepA* (GenBank Y10480.1 and X86702.1) showed 99 %ID to their alignment to pBFO18_1 (not shown) (51,52). Strain S01 putative plasmid sequence pBFS01_2 (8,331 bp) showed 99.87 %ID to the 1486bp partial sequence of *B. fragilis* plasmid pBF388c (GenBank AM042593.1), a 8.3kb conjugative plasmid harbouring *nimE* and IS*Bf6* (53).

None of the three putative plasmid sequences of strain BFO18 could be identified using the PLSDB but querying the NCBI nucleotide database using BLASTn revealed hits for all three. The hits corresponded to circularised sequences (%ID: 99.56-99.96, %COV: 100) assembled from mobilome metagenomic sequencing of the uncultured caecum content from a rat trapped at Bispebjerg Hospital in Copenhagen, Denmark (two hours’ drive from Odense University Hospital where BFO18 was isolated from a patient’s blood culture) (Supplementary Table S3) (47,54). BLASTn searches of the remaining unidentified putative plasmids from the other strains did not reveal complete hits.

Using ABRicate with the plasmid replication domains collected from the Pfam database, all putative plasmids, except pBF017_1 and pBFS01_1, were found to have recognised replicon domains (Table 4). DNA fragments of sizes matching pBFO17_1 and pBFS01_1 were detected by PFGE of S1 endonuclease restriction enzyme treated plasmid DNA extracts (Supplementary Figure S4) and the circular structures of the two sequences lacking a predicted replication domain, were confirmed manually by visually inspecting BLASTn mapping of ONT sequences longer than 10 kbp to the assembled plasmid sequences with CLC Genomics Workbench 10 (Qiagen). Eleven and 22 ONT reads spanned the complete lengths of pBFO17_1 and pBFS01_1 respectively and contained no other elements. pBFO17_1 and pBFS01_1 demonstrate a degree of similarity of close to 100%, except for an approximate total of 7,500 bp transposase and prophage sequences in pBF017_1 (Figure 3). No alignment to chromosomal sequences of any of the included *B. fragilis* isolates was observed using progressiveMauve (not shown) (55).

**Figure 3.**
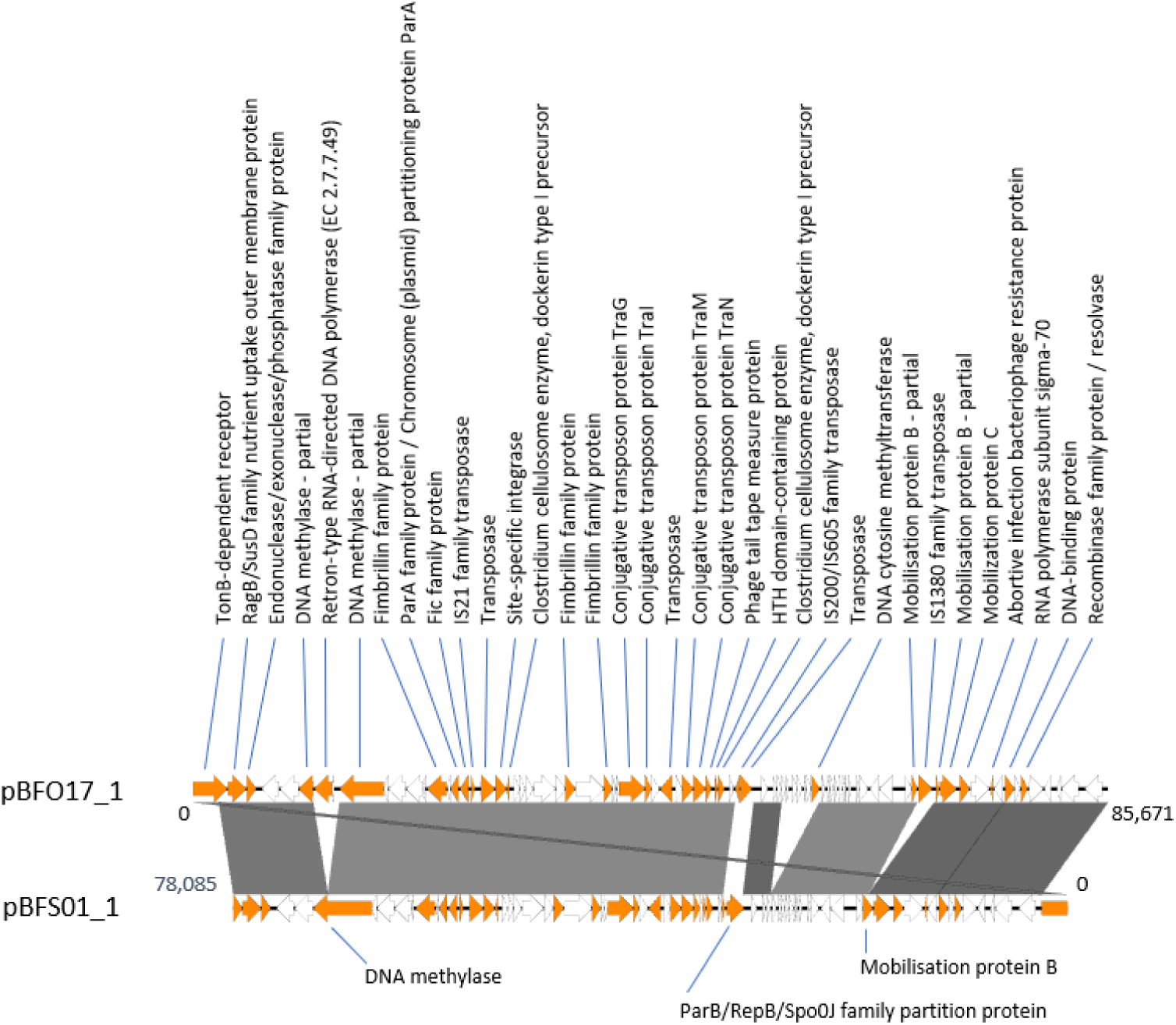
Linear representation of an alignment of putative circular plasmid sequences pBFO17_1 pBFS01_1. Comparison of the putative circular plasmids pBFO17_1 and pBFS01_1 (reverse complement for better visualisation) using EasyFig (82). EasyFig uses BLAST to identify sequences of similarity. Sequence similarities of >98% is indicated by full colouring, a darker colour indicates a higher %ID. Products of annotated CDS’ are shown. CDS’ annotated as hypothetical or Domain of Unknown Function are coloured white. The two sequences show a very high degree of similarity. pBFO17_1 is 7,586 bp longer than pBFS01_1. This is mainly due to the insertion of a reverse transcriptase (pBFO17_1, 11367..13034) (disrupting a DNA methylase), the insertion of prophage from position 56125 to 61162) (identified as an incomplete prophage using PHASTER (83)) and an IS*1380* family-like transposase (67933..69237). The regions pBFO17_1 50711..52501 and pBFS01_1 32248..30304 are not similar. Possibly, the insertion of two transposases in pBFO17_1 have excised most of the ParB-family DNA partitioning protein in the corresponding sequence range in pBFS01_1.

The GC content of pBFO17_1 and pBFS01_1 are 36.78% and 36.04% respectively. These lie within the range for the *Bacteroides* genus but differ from the expected value for *B. fragilis* (43%), which could indicate that the putative plasmids do not originate from *B. fragilis* (56). After supplementing the PGAP annotations with RAST annotation (57), 63% (pBFO17_1) and 59% (pBFSO1_1) of CDS’ remained annotated as hypothetical or as domain of unknown function. Of the annotated CDS’ the majority were associated with mobilisable features, plasmids and phages such as *parA* and *parB*, DNA partitioning proteins, conjugative transposon proteins, transposases, DNA binding motif domain containing proteins, and reverse transcriptase protein. The results above support the assembly data suggesting these two sequences are in fact plasmids.

### Detection of antimicrobial resistance genes and Insertion sequence elements

We used ABRicate to screen assemblies for AMR genes (ResFinder, NCBI and CARD databases supplemented with sequences for *bexA* and *bexB*) and IS elements (IS-finder database); several AMR genes, possible homologs to known AMR genes and IS elements adjunct to the AMR genes were detected (Table 5). Of note, isolate BFO17 contains two homologs of the metronidazole resistance gene *nimJ* (with a 100% consensus) and two isolates, S01 and BFO85, harbour two homologs of the tetracycline resistance gene *tetQ*. Homologs to *bexA* and *bexB* were identified with 73.53-99.12 %ID and were all confirmed with BLASTx searches against the NCBI nr database, as was done in our previous study (14). Partial hits for *ugd* was observed for several isolates, but with low %ID and %COV, and possible represent identification of conserved domains, but not *ugd* homologs. Increased expression of the *cfiA* metallo-beta-lactamase gene, *nim*-family 5-nitroimidazole genes and *erm* genes is partly regulated through IS elements containing promoter sequences. Full length IS elements could be identified upstream of 11 (79%) of 14 *cfiA, nim* and *erm* genes and upstream of two of three *CfxA4* genes and the *OXA-347* gene identified in BFO42. The described *Bacteroides fragilis* promotors TAnnTTTG (−7) and TG or TTG or TGTG (−33) (58) were searched for manually but could not be identified upstream of the two *cfiA* genes in isolates BFO67 and BFO85 or the *ermB* gene in BFO85 for which no IS elements could be detected upstream (not shown).

**Table 5.**
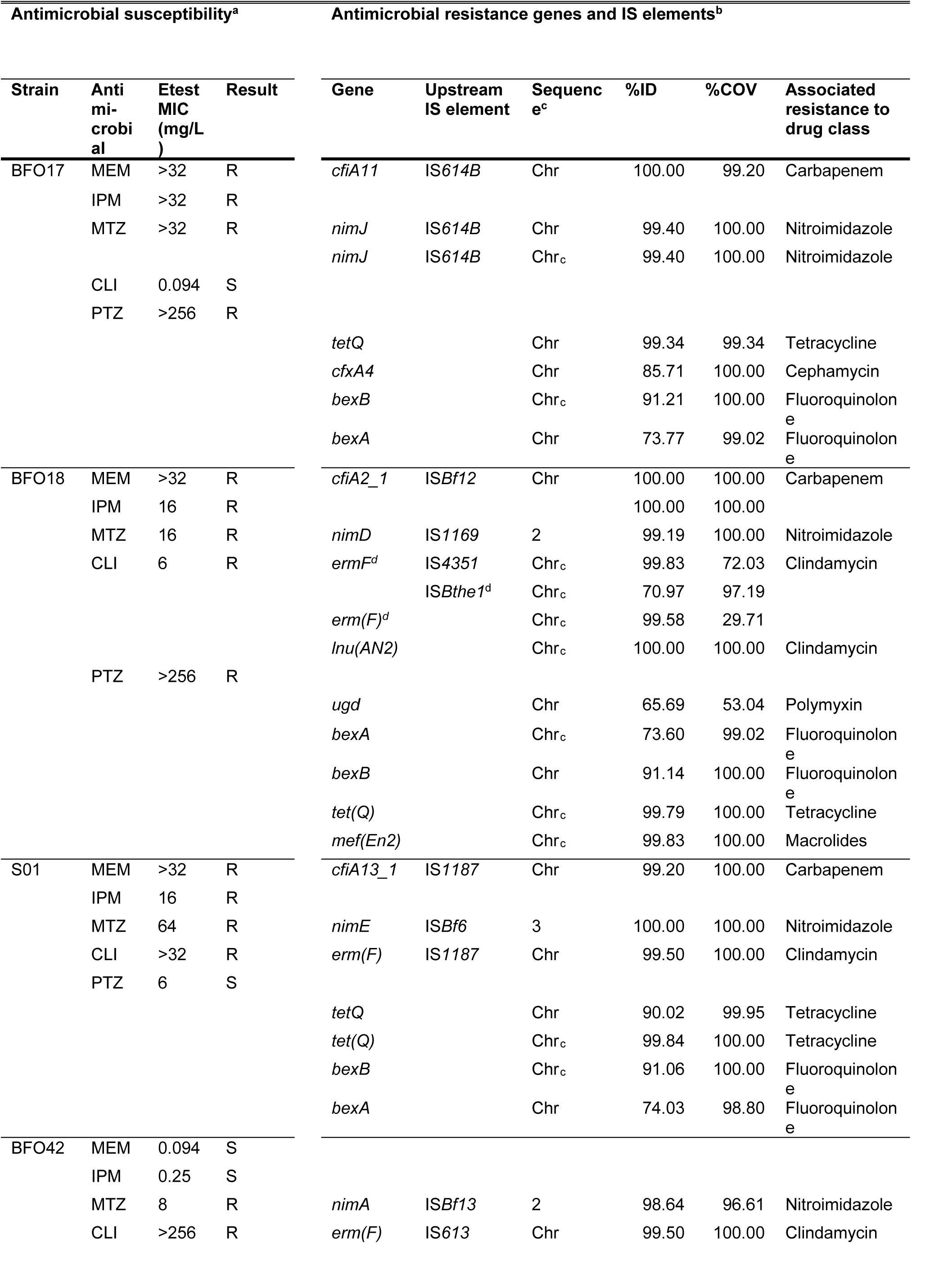
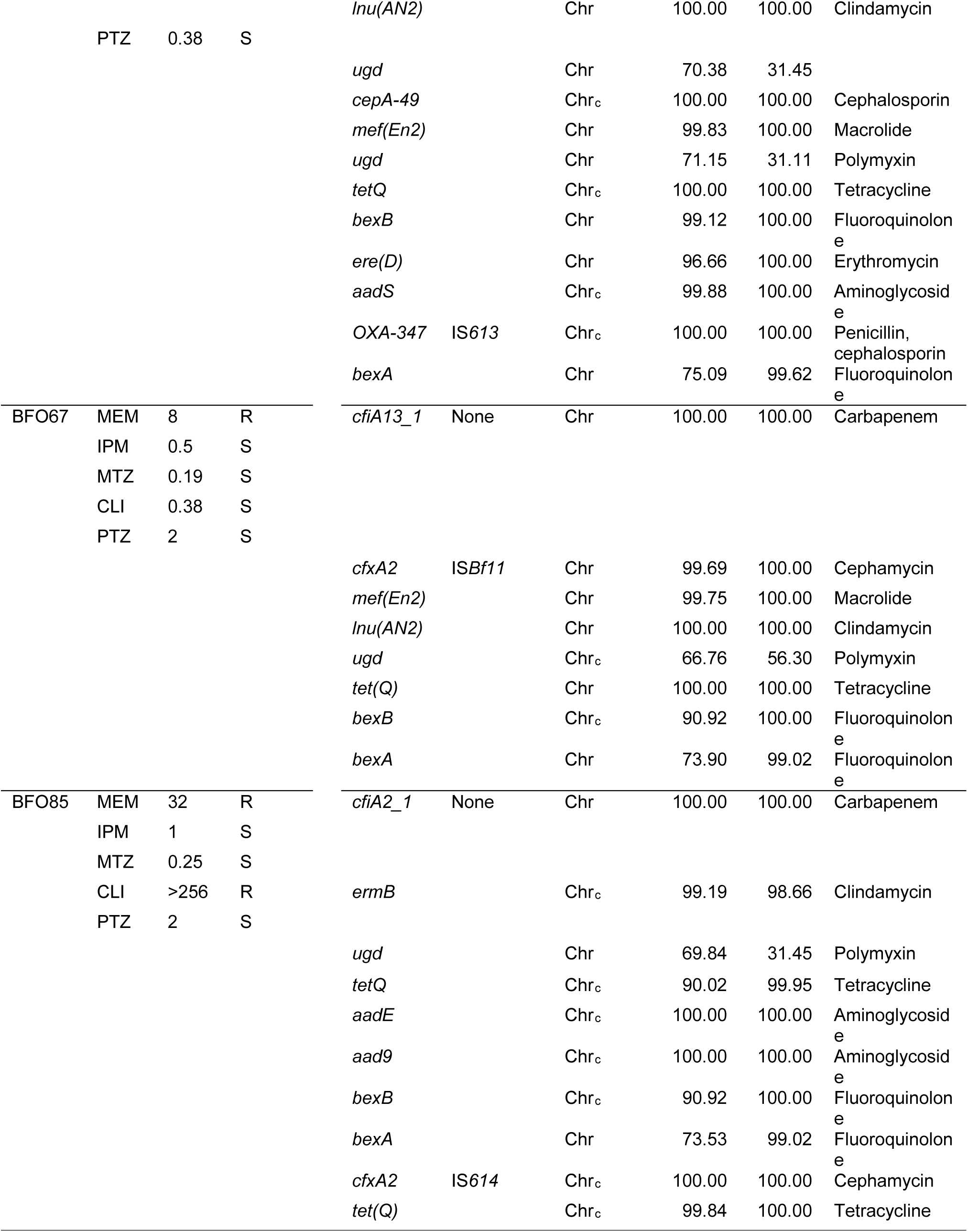
Antimicrobial susceptibility and resistance genes and IS elements for the six MDR *B. fragilis* strains. Identified genes are displayed next to the relevant antimicrobials. Identified IS elements in correct orientation (opposite strand) directly upstream of the genes are included. The %ID and %COV refer to the gene hit. Hits with %ID or %COV <98% were confirmed with BLASTx searches. The hits for *ugd* represent possible homologs for genes coding for PmrE, which is involved in polymyxin resistance in Gram-negative bacteria. Full ABRicate results with nucleotide positions and information on the IS elements is available the Supplementary Tables S4 AMR-IS-results.xlsx. Notes: ^a^ Results from previously published work following EUCAST breakpoints (14). ^c^ A _c_ denotes complement strand. ^d^ A transposase has inserted itself, splitting the *ermF* gene in two. Abbreviations: %ID; percent identity, %COV; coverage percentage, Chr; chromosome.

### Correlation between identified genes and IS elements and phenotypical resistance

As in our previous study, the *cfiA* gene was identified in the five meropenem resistant isolates (Table 5). All the *cfiA* genes were found on the chromosomal sequences. Complete IS elements were identified upstream of the *cfiA* genes in BFO17, BFO18 and S01, but not in BFO67 or BFO85. MICs for meropenem and imipenem were lower for these two isolates. *Nim* genes (−*A, -D, -E* and -*J*) could be found in the four metronidazole resistant isolates, all with complete IS elements upstream. Three of the *nim* genes were found on putative plasmids of the respective isolates. The four clindamycin-resistant isolates all carried *erm*-genes with upstream IS elements. A transposase was inserted in the *ermF-gene* in isolate BFO18, splitting it in two and the same isolate demonstrated a lower clindamycin MIC (6 mg/L) than the other three clindamycin resistant isolates.

## DISCUSSION

### Hybrid genome assembly produces high quality *B. fragilis* genomes

The primary aim of this study was to select and validate an assembly method to reliably complete chromosome and plasmid assembly of *B. fragilis* genomes. From 141 assembly variations, a hybrid approach using Filtlong filtered and Canu corrected ONT reads with quality filtered Illumina reads as input to Unicycler produced a complete, closed assembly of *B. fragilis* CCUG4856T with high similarity to the reference assembly of the original Sanger sequenced reference assembly. An 88kb inversion was observed when comparing the two assemblies. Cerdeño-Tárraga and colleagues observed difficulties in resolving certain regions of the Sanger sequenced assembly of NCTC9343 due to invertible regions with flanking inverted repeat sequences (21). The observed inversion in the hybrid Unicycler assembly, could be due to a) a superior assembly where the longer ONT reads have overcome the shortcomings of the shorter Sanger sequences, b) an incorrect assembly by Unicycler, c) a biological difference that has occurred over time between the strain stored at NCTC and CCUG, or d) a biological difference that occurred during the culturing of the strain, with dominance of a clone with the inversion, prior to DNA extraction as part of this study. The observations that the inversion is also present in all the best assemblies from this study and assemblies from two other research institutions support the conclusions that the current hybrid Unicycler assembly represents the true orientation of the 88kb sequence.

### Complete genome assembly of three of the six multidrug resistant isolates required manual finishing

The assemblies of BFO18, S01, and BFO42 were completed by Unicycler without manual intervention, but the chromosomes of BFO17, BFO67, and BFO85 could only be closed by performing manual steps. The manual finishing steps are time consuming, difficult to replicate and are easily biased. In order to be implemented in routine clinical laboratories, large scale, automated, complete assembly of prokaryote genomes require robust methods with minimal human interaction. Genome assembly using another long-read assembler, Flye, supported the results of the manual finishing for two of three isolates. Flye is better at resolving repeats than miniasm, the long read assembler included in the Unicycler pipeline (59). One option could be to include the long-read assembly from Flye, in place of that of miniasm, to guide bridge building for the higher quality Illumina-only contigs produced in the first steps of Unicycler. To resolve repeats it is often necessary to have long reads that span the repeat. In prokaryotes repeats over 10kb are not unusual and they are often spanned by the ONT reads generated, even by novice researchers. But repeat regions of up to 120kb and duplications of 200kb have been described in some prokaryotes (17,18,60). ONT sequencing runs will routinely result in many reads that span the majority of repeats, but to obtain ONT reads that span specific 120-200kb repeats in a genome of interest still requires skill and a certain amount of luck. Protocols for ONT sequencing have been described that result in read lengths of over 2 Mb, but this requires skilled and experienced researchers and lab technicians and demands high amounts of very high quality input DNA and essentially sequencing of only one isolate per MinION flowcell (61).

ONT read depth did not serve as an indicator of whether the Unicycler assemblies would result in closed chromosomal contigs in this study. Final ONT read depth, prior to Filtlong filtering and Canu correction, ranged from 23-371x, but a high read depth alone, was not an indicator of closed contigs. The three assemblies BF017, BFO67, and BFO85 required manual finishing to complete the assemblies and had ONT raw read depths of 99-137x. After Filtlong filtering and Canu correction the median read lengths were 21,932-29,893b and read length N50 was 25,765-34,815b for the three isolates (Supplementary Table S1). Canu correction improved the Unicycler assembly of *B. fragilis* CCUG4856T by nearly all parameters. But whilst Canu correction of the data from the first sequencing run resulted in the complete assembly of BFO67, the assembly of S01 worsened slightly. Increasing the amount of ONT data for BFO67 fragmented the complete chromosome. However, increasing the ONT read depth did decrease the number of contigs per isolate in our study overall.

Defining an optimal approach for complete prokaryote genome assembly is a continuous process, as sequencing technologies and assembly software develop and mature. Ring and colleagues found that Canu correction prior to Unicycler hybrid assembly was superior to other hybrid assembly or long read assembly approaches for assembly of *Bordetella pertussis* genomes that contain long duplicated regions (18). Unicycler also performs well in other studies comparing assessing genome assemblers for bacterial genome and plasmid assembly (19). De Maio and colleagues recently published a preprint comparing hybrid assembly strategies for 20 *Enterobacteriaceae* isolates (20). In their dataset, simply randomly subsampling ONT reads to an approximate read depth of 100x was slightly superior to applying Canu correction or Filtlong filtering prior to Unicycler assembly. For 85% of isolates the expected number of circular contigs were all assembled. For only one additional isolate Canu correction or Filtlong filtering resulted in the assembly of the expected number of circular contigs. Manual steps, including down sampling ONT reads or removing the Canu correction are options to consider, if chromosomes are not complete and circularised after initial Unicycler assembly, providing ONT read depth of 100x is available.

We chose to benchmark a selection of widely used genome assemblers for short read, long-read and hybrid bacterial genome assembly as well as polishing tools for long read assemblies, but many other options have been published. Most assemblers and polishing tools were run using default parameters, and it is possible that further optimisation of settings for the individual software packages might have improved assemblies further than was demonstrated here. As sequencing technologies and assembly software continues to improve, continued validation of pipelines is advisable. Software such as poreTally provides user friendly options for benchmarking genome assembly pipelines prior to implementation (62).

### *Bacteroides* plasmids are not well represented in public databases

A secondary aim of this study was to identify plasmids in the hybrid assemblies. Automated tools have been developed and validated for identification of plasmids from genome assemblies or read data, but they are dependant of collated databases of known plasmid sequences. As such, tools such as PlasmidFinder or mlplasmids can be applied for plasmid identification for *Enterobacteriaceae* or *Enterococcus faecium*, but *B. fragilis* is not supported at the time of writing (63,64). Therefore, we evaluated putative plasmid sequences by sequence identity and length comparison using the PLSDB webpage, identifying plasmid replication domains, and using circularisation and relative coverage as indicators that a sequence represents a plasmid in a given isolate.

Only four of the twelve plasmid sequences from the seven isolates could be identified using the PLSDB and three of these were the same plasmid, pBFP35. Two other putative plasmids, pBFO18_1 and pBFS01_2 were likely plasmids pBF388c and pIP421 based on the partial sequences from these plasmids and plasmid length. This still leaves half of the circularised, putative plasmids unidentified. The two longer putative plasmids, pBFO17_1 and pBFS01_1, displayed a high degree of similarity, a GC% out of the normal range for *B. fragilis*, and a relative read depth of double the reads compared to the chromosome. Most annotated CDS’ were associated with mobilisable elements, but no known plasmid replication domains could be identified. From the sequencing data alone, we cannot conclude that they represent true plasmids, however the findings above and manual inspection of long read mapping support that inference.

There are only 14 complete plasmid sequences from cultured *Bacteroides* isolates in the PLSDB v2019_03_05, which is based on the NCBI RefSeq database. Many other *Bacteroides* plasmids have been partially described, and some are represented by partial sequences or marked as contig level in the NCBI nucleotide database (65–68). Metagenomic sequencing and genome assembly projects are expanding the public sequence databases and screening the NCBI nucleotide database, sequences with a high degree of similarity to the putative plasmid sequences from one patient isolate (BFO18) could be found. These originated from a rat caecum metagenomic plasmid sequencing project from Copenhagen, a few hours’ drive from Odense University Hospital. To understand and perform surveillance of the dissemination of plasmids there is a need for increased submissions of high quality, annotated and phenotypically validated sequences of bacterial isolates including plasmids. This study adds significantly to the number of complete plasmid sequences associated with *Bacteroides*.

### Complete assembly allows comprehensive identification of resistance determinants in *B. fragilis*

We also intended to comprehensively identify resistance genes and IS elements in the hybrid genome assemblies. Using ABRicate with several resistance gene databases and IS-element nucleotide sequences, the findings of our previous study were confirmed and enhanced. Assemblies from Illumina sequencing alone would only allow partial IS element identification (14). Now, with the complete assemblies, comprehensive identification of known IS elements upstream of the relevant resistance genes could be completed. In our first study we used ResFinder with the available database at that time. Now, by including several databases, and lowering the %ID threshold, the number of genes identified increased. Additionally, for as a result of the complete genome assembly of BFO17, we could now identify two copies of *nimJ*, while only one copy was identified in the short read draft assembly of the same isolate in the previous study. Husain and colleagues identified the presence of three copies of *nimJ* in strain HMW615, when describing the *nimJ* gene (69). We confirmed this finding by running ABRicate on the HMW615 assembly as done with the isolates of this study (not shown). Interestingly, RAST annotates a third *nim* gene (nucleotide positions 1,359,590..1,360,093) in the Unicycler hybrid assembly of BFO17, and the PGAP annotation includes an additional annotation of a pyridoxamine 5’-phosphate oxidase family gene (nucleotide positions 940,032..940,505), the family that includes the *nim*-genes. It is possible that one or more novel homologs of the *nim* are present in BFO17.

IS elements could be identified upstream of most relevant resistance genes. However, in three cases no IS element was present upstream of a resistance gene, even though the isolates displayed phenotypical resistance associated with increased expression of the specific gene. Known *B. fragilis* promoter sequences could not be identified upstream of the genes “missing” upstream IS elements, however *B. fragilis* promotors are still not completely described, so it is possible there are other unknown variants.

By selecting an optimal genome assembly strategy for *B. fragilis*, supplemented with minimal manual finishing efforts, and applying this to six multidrug resistant isolates, the number of complete *B. fragilis* genomes and plasmids in the public databases has now almost doubled. The future aim of performing antimicrobial resistance prediction based solely on WGS information for *B. fragilis* demands near-complete genomes for identification of IS elements upstream of resistance genes. However, we must caution that the absence of an IS element upstream of *cfiA* does not always correlate to susceptibility to carbapenems. Future studies are needed to address this, and utilising complete genome assembly for genome wide association studies is one approach that could be pursued. Technologies that provide a single solution for real-time, high-quality sequencing of long reads will be essential for implementing near real-time diagnostics of infectious diseases and characterisation of pathogens.

## ABBREVIATIONS

AMR: antimicrobial resistance;
WGS: whole genome sequencing;
IS: insertion sequence;
ONT: Oxford Nanopore Technologies;

## AUTHOR STATEMENTS

### Authors and contributors

The study was conceptualised by T.V.S. and U.S.J. Funding was secured by T.V.S., M.K., H.H. and U.S.J. Data curation and investigation was performed by T.V.S.. Formal analysis was done by T.V.S. and S.O-P.. Resources were provided by M.K., T.V.S., H.H. and U.S.J.. U.S.J, H.H. and M.K. supervised the work. T.V.S wrote the original draft and edited the manuscript. U.S.J., M.K., T.V.S., H.H., S.O-P. and H.M.W revised the manuscript.

### Ethical approval

Isolates were obtained as part of routine clinical care, and details about the isolates have previously been published. No ethical approvals were required.

## Acknowledgements

We are very grateful to Professor Henrik Westh, Department of Clinical Microbiology, Hvidovre Hospital, Denmark, for allowing use of their Computerome High Performance Computing cluster account for data analysis and to Mikala Wang, Dept. of Clinical Microbiology, Aarhus University hospital, for gifting us the MDR *B. fragilis* strain isolated at her department. We thank Valentina Galata, Saarland University, Saarbrücken, Germany (first author of the paper describing PLSDB) and Natalie Ring, University of Bath, Bath, United Kingdom (first author of reference (18)) for kind and helpful answers to questions via e-mail.

