## Supplementary Table S3 for "Complete genome assembly of clinical multidrug resistant *Bacteroides fragilis* isolates enables comprehensive identification of antimicrobial resistance genes and plasmids"

| Strain | Sequence | Length | Relative coverage | Best NCBI BLASTn hit | %COV | %ID | Length of accession |
|---|---|---|---|---|---|---|---|
| CCUG4 856T | Chr. | 5,205,133 | 1.00x | - | | | |
| | pBF9343 | 36,560 | 7.42x | - | | | |
| BFO17 | Chr. | 5,474,541 | 1.00x | - | | | |
| | pBFO17_1 | 85,671 | 1.85x | CP022754.1 | 11 | 86.06 | 5372666 |
| | pBFO17_2 | 5,594 | 23.03x | - | | | |
| BFO18 | Chr. | 5,302,644 | 1.00x | - | | | |
| | pBFO18_1 | 7,221 | 25.98x | LN852993.1 | 100 | 99.96 | 7160 |
| | pBFO18_2 | 4,137 | 50.80x | LN853919.1 | 100 | 99.77 | 4134 |
| | pBFO18_3 | 2,782 | 59.91x | HG796821.1 | 100 | 99.56 | 2784 |
| S01 | Chr. | 5,325,251 | 1.00x | - | | | |
| | pBFS01_1 | 78,085 | 2.29x | CR626928.1 | 0 | 98.74 | 36560 |
| | pBFS01_2 | 8,331 | 20.99x | CP002533.1 | 62 | 95.83 | 6277 |
| | pBFS01_3 | 5,595 | 22.67x | - | | | |
| BFO42 | Chr. | 5,141,257 | 1.00x | - | | | |
| | pBFO32_1 | 8,306 | 40.06x | KJ830768.1 | 70 | 97.35 | 11019 |
| | pBFO32_2 | 5,594 | 40.15x | - | | | |
| BFO67 | Chr. | 5,478,614 | 1.00x | - | | | |
| | pBFO67_1 | 6,129 | 94.15x | AB646744.1 | 26 | 79.49 | 12817 |
| BFO85 | Chr. | 5,504,076 | 1.00x | - | | | |

Supplementary Table S3 – BLASTn results for putative plasmids not identified with PLSDB. Putative plasmid sequences not identified with PLSDB were BLASTn'ed against NCBI nr database (prokaryotes). The top hit is listed in the table above with NCBI accession numbers. Sequences similar to all the putative plasmids in BFO18 were identified in the NCBI nr database. All originating from plasmid centric metagenomic sequencing and assembly of the contents of the caecum of a rat trapped at Bispebjerg Hospital, Copenhagen, Denmark (1). %COV: Coverage percent, %ID: Percent identity.
