## Supplementary figures for "Complete genome assembly of clinical multidrug resistant *Bacteroides fragilis* isolates enables comprehensive identification of antimicrobial resistance genes and plasmids"

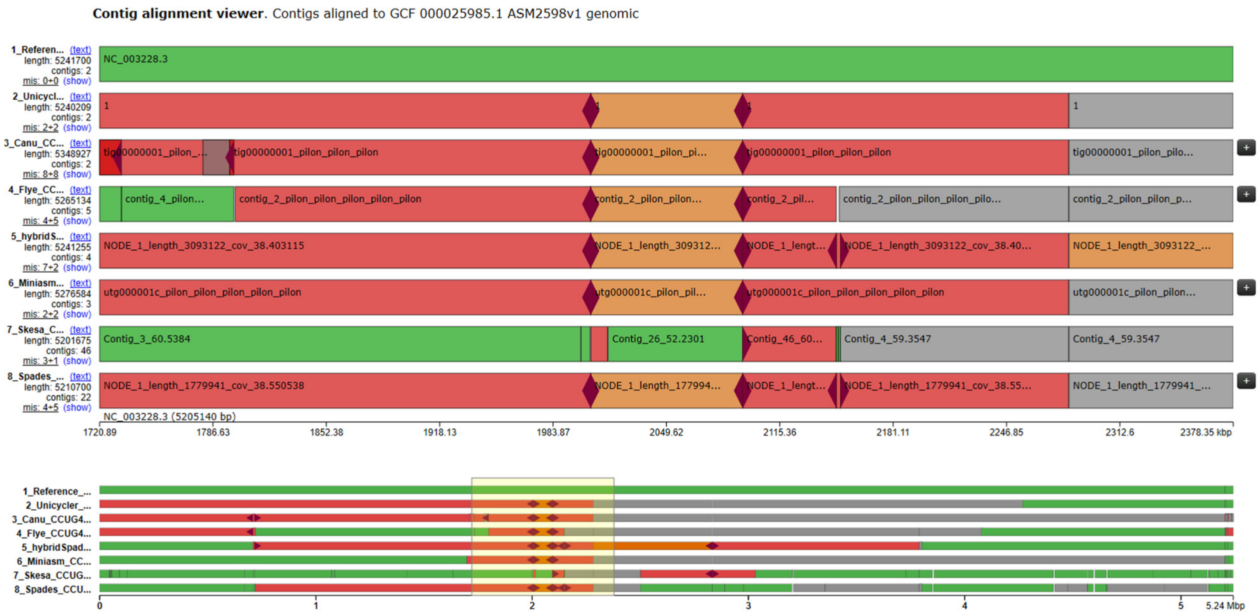


Supplementary Figure S1. Alignment of best assemblies per assembler. Visualisation of contig alignments generated by Quast of the best assembly per assembler to the reference genome assembly GCF_000025985.1 (top) using the Icarus viewer (1). Assemblies are numbered 1-8 to order the alignments in the figure. The 88,045 bp inversion is present in all assemblies, including those from ONT reads (Canu, Flye and Miniasm) or Illumina reads only (Skesa and Spades). Borders of the inversion are indicated with the dark grey diamond shape in the Icarus view. In contrast to the Spades assembly, where the inversion is included in the contig NODE_1, the Skesa assembler has produced a more fragmented assembly. Skesa contig_26 spans most of the 88,045 bp inversion and is not part of a longer contig. Skesa uses conservative heuristics and is designed to create breaks in repeat regions in the genome.

Supplementary figures

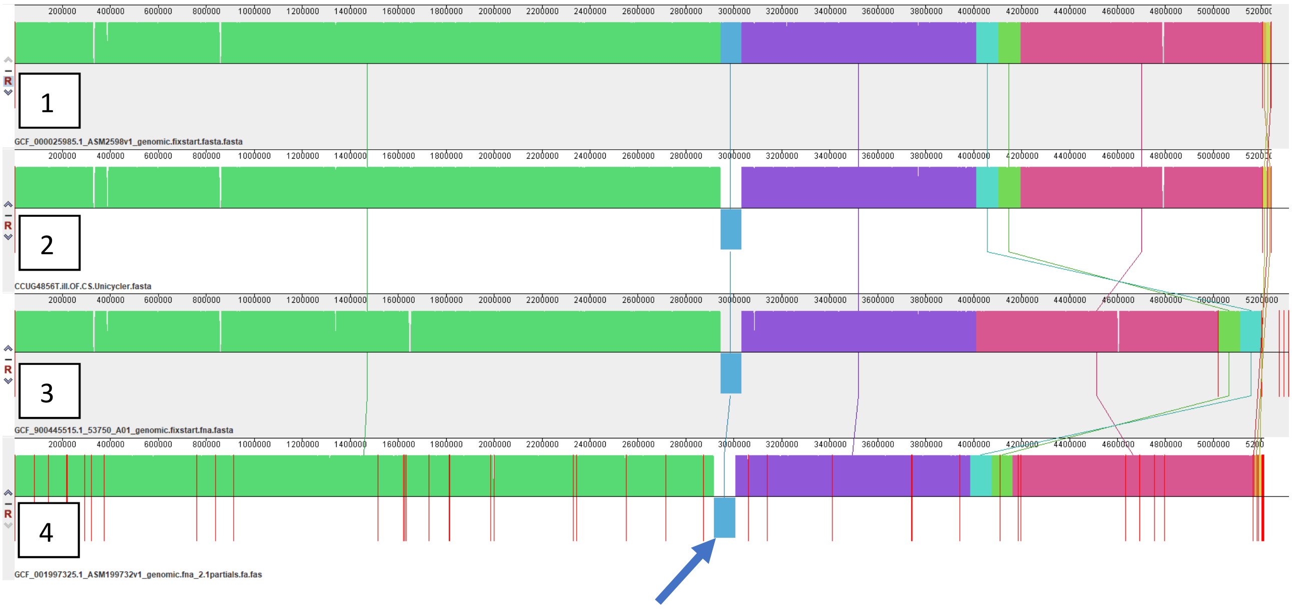


Supplementary Figure S2. Alligment of available assemblies of *B.fragilis* CCUG4856. Alignment using progressiveMauve (2) of available NCBI RefSeq assemblies of *B. fragilis* CCUG4856T(=ATCC25285=NCTC9343) as well as the best Unicycler hybrid assembly from this study. Assemblies 1: NCTC9343 Sanger sequenced, two contigs (NCBI RefSeq GCF_000025985.1); 2: CCUC4648T Hybrid assembly using Unicycler with ONT and Illumina reads, two contigs (this study); 3: NCTC9343 assembly from PacBio data by the Sanger Institute as part of the NCTC 3000 project (http://www.sanger.ac.uk/resources/downloads/bacteria/nctc/), five contigs (NCBI RefSeq GCF_900445515.1); 4: ATCC25285 assembled using CLCbio Genome Workbech with Illumina Miseq reads submitted by the California Department of Health, 59 contigs (NCBI RefSeq GCF_001997325.1). Assembly 1 and 3 were rotated to start at the *dnaA* gene using Circlator's fixstart and contigs for no. 4 were reordered using Mauve for better visualisation. NCBI genome assembly GCA_900227845.1 is labelled *B. fragilis* NCTC9343, however, alignment and kmer-analysis (using KmerFinder 3.1 (https://cge.cbs.dtu.dk/services/KmerFinder/) (3) clearly shows that the accession is not *B. fragilis*, but rather an *Enterobacteriaceae.* The accession is therefore not included in this figure. The 88kb inversion observed in all the best assemblies of this study is also present in the PacBio and Illumina assemblies submitted by unaffiliated researchers (marked with an arrow). The assembly from the NCTC 3000 project is missing plasmid pBF9343 and has additional sequences that are not found in the original Sanger sequenced assembly or the hybrid assembly from this study.

Supplementary figures

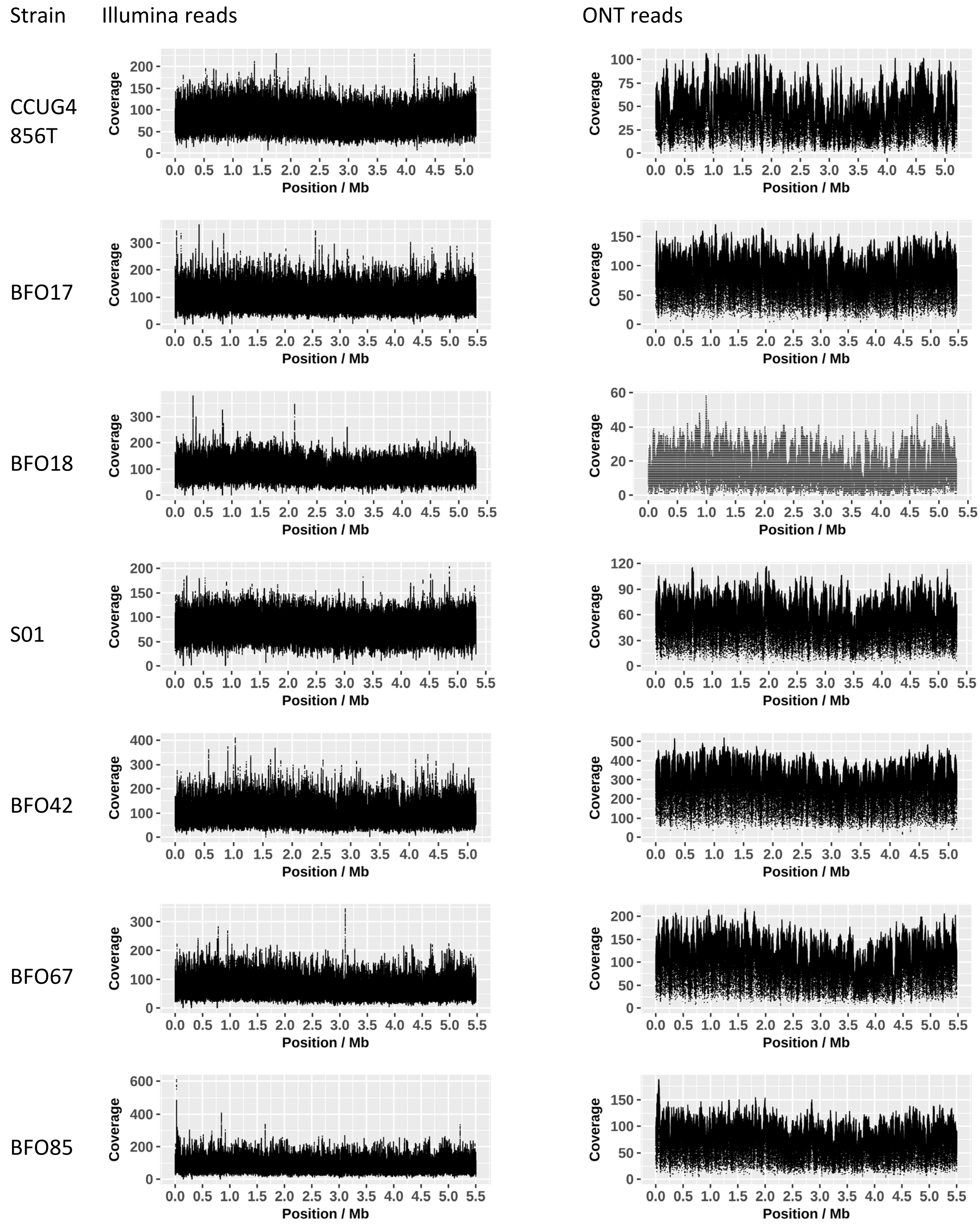


Supplementary Figure S3 Alignment of Illumina and ONT reads to the chromosomes of the respective strains. Quality filtered and trimmed Illumine reads were aligned using BWA MEM and Filtlong filtered Nanopore reads were aligned using minimap2. Coverage calculation was performed using samtools depth. The coverage was consistent across most genomes. In BFO85 both Illumina and Nanopore coverage is almost twice as high for a region at approximately 25kb-38kb, suggesting this is not a bias of a sequencing technology. The 13kb region does not contain annotated mobilisation proteins. But the presence of a gene predicted to encode RNA polymerase sigma-70 factor (29340..29897). This could represent an approximately 13kb repeat that has not been resolved in this assembly.

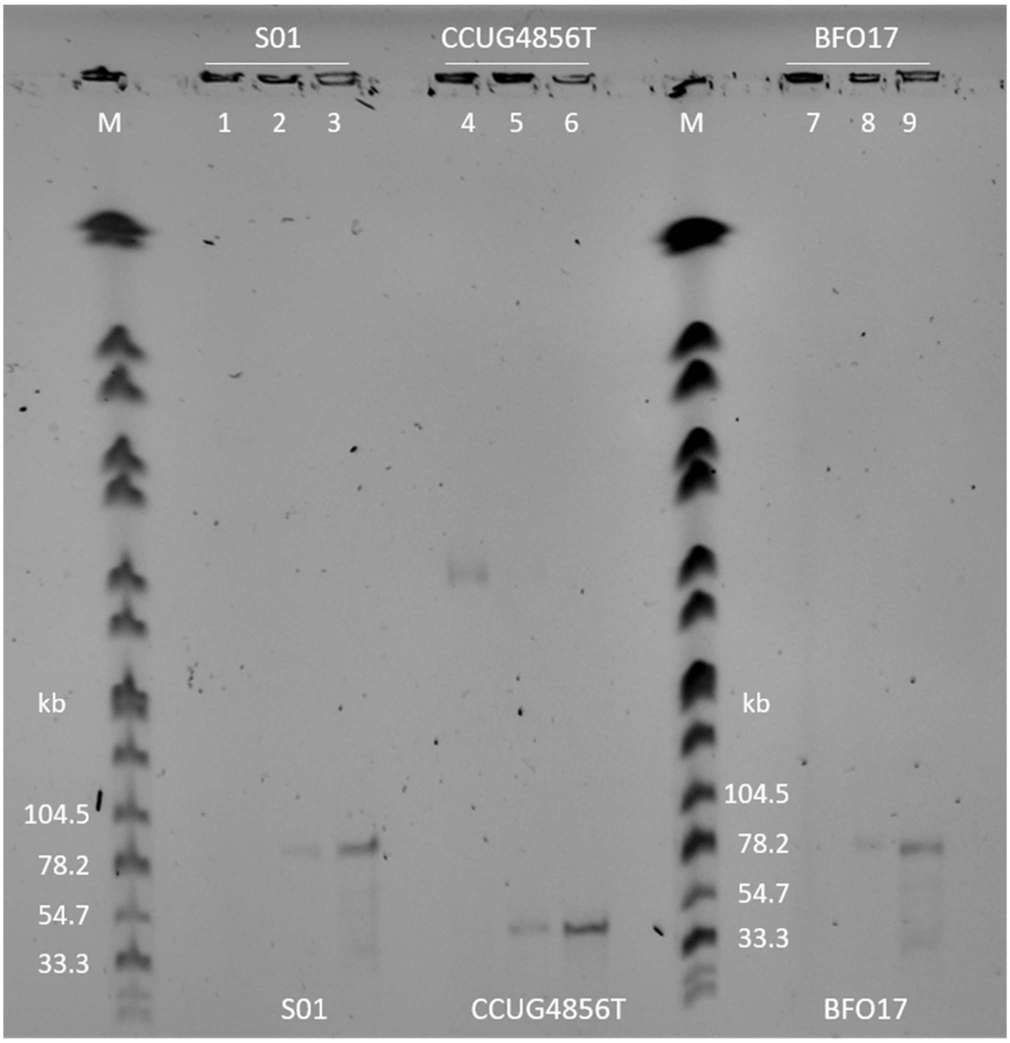


Supplementary Figure S4. PFGE patterns of S1 endonuclease digested plasmid preparations of *B. fragilis* strains S01, CCUG4856T, and BFO17. Lanes 1-3; Strain S01 untreated, treated with 5U S1, and 100U S1. Lanes 4-6; Strain CCUG4856T untreated, treated with 5U S1, and 100U S1. Lanes 7-9; Strain S01 untreated, treated with 5U S1, and 100U S1. Lanes M; DNA size marker (*S*. Braenderup H9812 digest with *Xba*I). Plasmid DNA was extracted using Plasmid Mini kit (Qiagen). S1 digestion: Plasmid DNA containing agarose plugs were added to 40 µL 5X S1 Reaction Buffer (ThermoFisher Scientific) and 5 or 100 units S1 Nuclease (ThermoFisher Scientific) and sterile water to a total volume of 200 µL. Reactions were incubated at room temperature for 30 minutes before being subjected to PFGE (6.8 – 38.4 sec, 19 hours).
